# A pathogen α-L-arabinofuranosidase degrades host-derived immunogenic oligosaccharides to suppress plant immunity

**DOI:** 10.64898/2026.08.18.742768

**Authors:** Cristian Carrasco-López, Diego Rebaque, Felipe de Salas, Sergio López-Cobos, Inés Vegas-Lorenzo, Gonzalo Vílchez-Pinto, María Garrido-Arandia, María Jesús Martínez, Hugo Mélida, Antonio Molina, Andrea Sánchez-Vallet

**Affiliations:** Centro de Biotecnología y Genómica de Plantas, Universidad Politécnica de Madrid (UPM) - Instituto Nacional de Investigación y Tecnología Agraria y Alimentaria/Consejo Superior de Investigaciones Científicas (INIA/CSIC), Campus de Montegancedo UPM, 28223, Pozuelo de Alarcón, Madrid, Spain; Departamento de Ingeniería y Ciencias Agrarias, Facultad de Ciencias Biológicas y Ambientales, Universidad de León, 24071, León, Spain; Instituto de Biología Molecular, Genómica y Proteómica (INBIOMIC), Universidad de León, Campus de Vegazana, 24071, León, Spain; Centro de Investigaciones Biológicas Margarita Salas, Spanish National Research Council, C/ Ramiro de Maeztu 9, 28040 Madrid, Spain; Departamento de Biotecnología-Biología Vegetal, Escuela Técnica Superior de Ingeniería Agronómica, Alimentaria y de Biosistemas, Universidad Politécnica de Madrid (UPM), 28040, Madrid, Spain

**Keywords:** Plant cell wall, carbohydrate active enzymes (CAZymes), pathogen effector, cell wall signaling, Damage-associated molecular pattern (DAMP), Septoria tritici blotch (STB), *Triticum aestivum*

## Abstract

Plant cell wall fragments released during pathogen attack can act as signalling molecules that trigger immune responses. Successful pathogens have potentially evolved diverse strategies to evade host recognition, including minimizing the accumulation of cell wall-derived elicitors. However, the mechanisms underlying this process remain largely unknown. Here, we characterized *Zt*GH54, an α-L-arabinofuranosidase from the wheat pathogen *Zymoseptoria tritici*, that is essential for the acquisition of sugar nutrients from arabinan and arabinoxylan wall polysaccharides. *Zt*GH54 also hydrolyzes immunogenic oligosaccharides derived from arabinoxylan to prevent host recognition. Remarkably, this strategy is effective only in a subset of wheat cultivars, as the contribution of *Zt*GH54 to virulence is cultivar-dependent. While a *Zt*GH54 substrate is broadly recognized in wheat, one of the products generated by *Zt*GH54, xylotetraose, is recognized only by specific wheat cultivars, revealing natural variation in the perception of xylan-derived oligosaccharides. These findings establish *Zt*GH54 as a key virulence factor that simultaneously exploits cell wall host resources and suppresses wheat immunity through the precise hydrolysis of plant cell wall-derived signals.

## INTRODUCTION

Pathogens engage in intimate interactions with their plant hosts. Pathogens damage their host tissues to grow and acquire nutrients, while plants activate defense responses to prevent pathogen colonization. Thus, pathogen progression is strongly dependent on their capacity to attack plants while remaining undetected. This challenge is particularly evident at the plant cell walls (*1–3*).

Plant cell walls constitute a complex and highly dynamic structure. They act as structural barriers that prevent the access of pathogens, and are a source of nutrients for plant co-existing microorganisms (*4*). Thus, pathogens have developed a highly adapted repertoire of enzymes targeting distinct cell wall components, such as cellulose, pectins and hemicelluloses (*5–10*). Several of these cell wall modifying enzymes (CWMEs) hydrolyze glycosidic linkages of plant cell wall polysaccharides to acquire carbohydrate nutrients and grow in the apoplast (*2*). However, plants have evolved sophisticated monitoring systems to recognize plant cell wall damage and trigger an immune response (*4*, *11*, *12*). One of these surveillance mechanisms involves the recognition of cell wall-derived oligosaccharides released during pathogen attack. These recognized cell wall-derived oligosaccharides are classified as damage-associated molecular patterns (DAMPs) and are perceived by plasma membrane-localized pattern recognition receptors (PRRs), activating immune responses that include calcium influx, accumulation of reactive oxygen species (ROS), and stomatal closure (*11*, *13*). This recognition is generally considered to be conserved across genotypes within a plant species (*2*, *11*, *14*). To counteract this defense response, pathogens use diverse strategies. For example, pathogens tightly regulate the expression of CWMEs to produce them only at specific stages of plant infection (*15–18*). This tight regulation has been shown to be important to prevent early recognition of the fungus by the host (*19*). Modification or hydrolysis of cell wall-derived elicitors (DAMPs) is an additional strategy used by host-associated microorganisms to prevent host recognition (*2*). The root-colonizing fungus *Serendipita indica* co-expresses an endo-xylanase, an acetyl-xylan esterase, and an exo-xylanase to hydrolyze xylan and degrade the released immunogenic oligosaccharides, preventing host recognition and facilitating colonization of barley (*20*). Similarly, the bacterial pathogen *Ralstonia solanacearum* secretes a polygalacturonase to break down immunogenic pectin-derived oligogalacturonic acid fragments into galacturonic acid residues, thereby dampening host immunity and capturing nutrients within the tomato xylem (*21*). Beyond these examples, the mechanisms used by pathogens to evade host recognition and obtain essential nutrients during plant infection remain mostly unknown.

Plant cell wall composition and structures vary widely among plant taxa (*9, 22*, *23*). Pathogens harbor specialized repertoires of CWMEs adapted to degrade the cell walls of their specific host species (*24–26*). In commelinid monocots, such as wheat (*Triticum aestivum*), arabinoxylan (AX) is a major hemicellulose (*27–29*). This polymer consists of a linear 1,4-β-ᴅ-xylan backbone with terminal α-L-arabinofuranosyl side chains at the *O*-2 and *O*-3 positions of the xylopyranosyl units (*30*). Hydrolysis of AX requires the activity of endo-1,4-β-ᴅ-xylanases as well as exo-acting enzymes, including α-L-arabinofuranosidases that release the terminal α-L-arabinofuranosyl residues linked to the xylopyranosyl units (*31*). Several pathogen endoxylanases have been described to mediate plant invasion (*32–34*) and one arabinofuranosidase contributes to *Magnaporthe oryzae* virulence (*35*). In *Z. tritici* an α-L-arabinofuranosidase from the family GH54 was associated with virulence and cultivar-specificity in wheat (*36*). Although these enzymes are required for pathogen progression, their activity can generate xylotetraose (Xyl4) and 3^3^-α-L-arabinofuranosyl-xylotetraose (XA^3^XX), which have been characterized as DAMPs (*37–39*). Thus, pathogens face the challenge of degrading cell wall components while limiting the production of cell wall-derived elicitors.

Apoplastic pathogens grow in tight contact with plant cell walls and produce CWMEs to breach them and colonize the host. For example, *Zymoseptoria tritici* penetrates through the stomata, grows exclusively in the wheat apoplast during infection, and produces tightly regulated CWMEs (*3*, *15*, *16*, *18*). The best-characterized CWME of *Z. tritici* is *Zt*GH45, an endoglucanase that releases mixed-linkage glucan (MLG)-derived oligosaccharides that trigger plant immune responses. *Zt*GH45 is produced at late stages of infection to prevent early recognition by the host and facilitate colonization (*19*). Despite the apparent importance of cell walls for *Z. tritici* colonization, the mechanisms by which this pathogen hydrolyzes plant cell walls at earlier stages of infection to acquire nutrients while avoiding detection by the host remain largely unknown.

In this study, we characterized *Zt*GH54, an α-L-arabinofuranosidase from *Z. tritici,* and demonstrated that it plays a dual role in virulence by facilitating nutrient acquisition and suppressing wheat immunity. Despite its positive contribution to virulence, this effect is observed only in a subset of wheat cultivars, as some cultivars recognize *Zt*GH54 and the products of its enzymatic activity. This cultivar-specific recognition highlights a molecular arms race at the plant cell wall, driving the co-evolution of pathogen CWMEs and wheat immune recognition.

## RESULTS

### The cell wall-modifying gene *ZtGH54* is strongly induced during *Z. tritici* infection

We conducted a genome-wide search using the dbCAN3 database (*40*) to identify *Z. tritici* CWMEs involved in hydrolyzing wheat cell walls. Since arabinoxylan constitutes the major polysaccharide in the hemicellulosic fraction of commelinid monocot cell walls (*27–29*), we focused on *Z. tritici* genes encoding secreted CWMEs with predicted catalytic activity against arabinan and xylan. This *in silico* process identified 11 candidate genes from various functional families, including GH3, GH11, GH43, GH51, GH54, GH62, GH115, and CE5 (Fig. 1A), with only *Zt*GH54 predicted to have activity against both substrates. To assess the contribution of these predicted CWMEs to plant colonization, we assessed whether they were expressed during wheat infection, using previously published transcriptomic datasets collected when no symptoms were developed (7 dpi), when the first symptoms appeared (12 dpi), and during the necrotrophic phase (14 and 28 dpi; *15*, *41*). All the identified potential xylanases and arabinanases exhibited very low expression levels under *in vitro* conditions. During plant infection, they were expressed at different time points. For example, the expression of the genes *Zt3D7_G6975* (an endo-1,4-β-D-xylanase from family GH11) and *Zt3D7_G6976* (an acetyl xylan esterase from family CE5) was induced at 14 and 28 days post infection (dpi), while the predicted exo-acting β-xylosidase *Zt3D7_G5219* (family GH3) reached its highest expression levels at 28 dpi. In contrast, the *Zt3D7_G4448*/*ZtGH54* gene, which encodes a putative α-L-arabinofuranosidase, belonging to the GH54 family, reached its maximum levels of expression at 12 dpi. Its expression levels decreased at the later stages of infection, with minimal transcript accumulation once the wheat leaves are completely necrotic. A similar expression pattern was observed for *Zt3D7_G9244* (an α-L-arabinofuranosidase from family GH43), although its expression levels were significantly lower than those of *Zt*GH54 (Fig. 1A). The high expression of *ZtGH54* at the early stages of infection suggested its potential role in plant infection.

**FIGURE 1.**
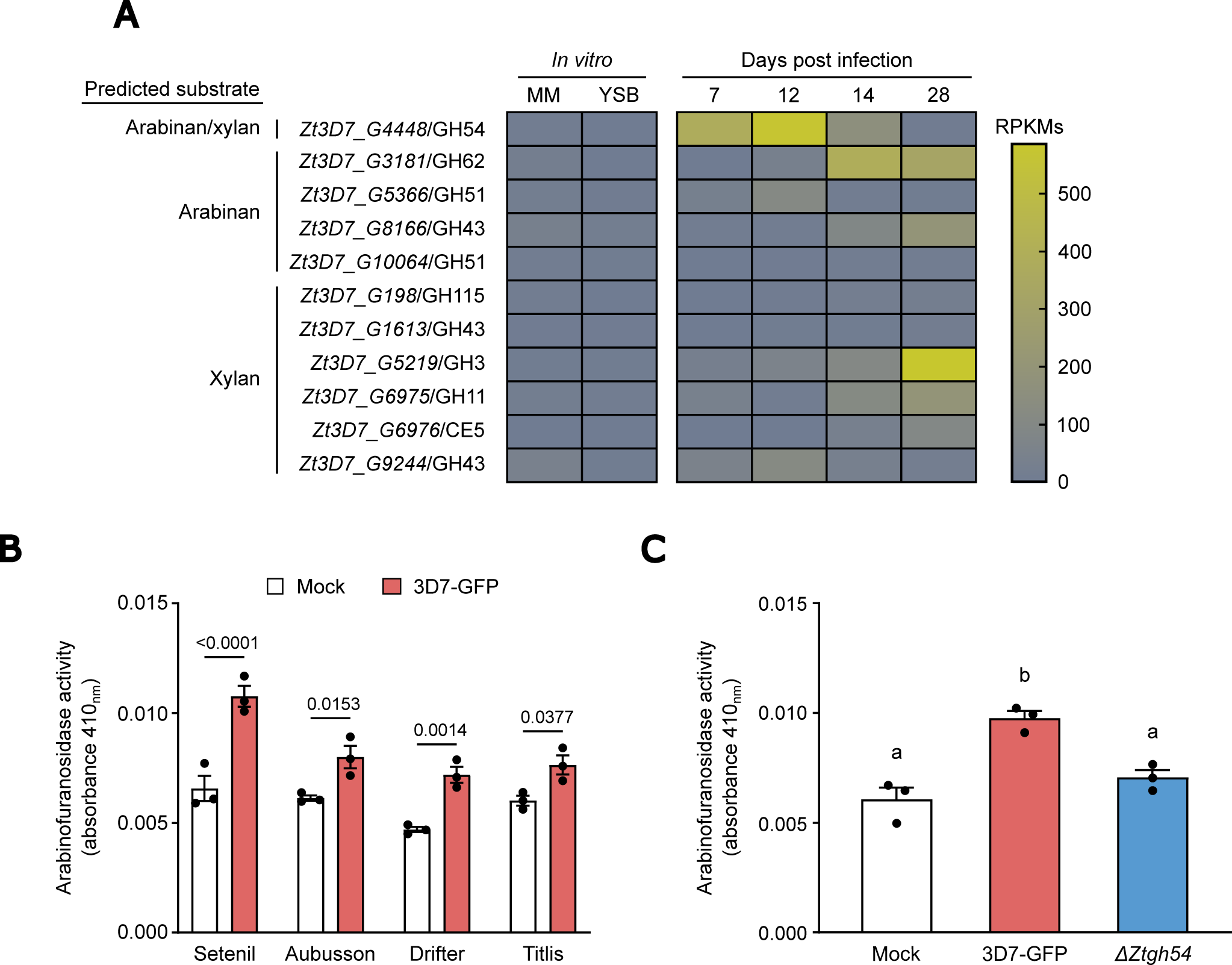
Induced arabinofuranosidase activity during wheat infection is primarily mediated by *Zt*GH54. **A.** Expression levels (Reads per kilobase of transcript per million reads mapped; RPKM) of *Z. tritici* genes encoding secreted CWMEs with predicted catalytic activity against arabinan, xylan, or both substrates in the 3D7 strain on minimal medium (MM) and yeast extract sucrose broth (YSB); and during wheat infection at 7, 12, 14, and 28 days post infection (dpi). Data were obtained from previously published RNA-seq studies (NCBI accessions: SRP152081 and SRA SRP077418). **B.** Apoplastic arabinofuranosidase activity in cultivars Setenil, Aubusson, Drifter, and Titlis, non-infected (mock, white bar) and infected with 3D7-GFP (control, red bar) at 7 dpi using the substrate *p*-nitrophenyl-α-L-arabinofuranoside (O-PNPAF) and determining the absorbance (410_nm_) of the arabinofuranosidase activity product. *P* values according to two-way ANOVA followed by Šídák test between plants infected with the 3D7-GFP strain and mock-treated plants are displayed in the plot. **C.** Apoplastic arabinofuranosidase activity (absorbance at 410_nm_) at 7 dpi in cultivar Setenil non-infected (mock) or infected with 3D7-GFP or a deletion mutant of *ZtGH54* (Δ*Ztgh54*) using the substrate O-PNPAF. Different letters indicate significant differences (*P* value <0.05) according to one-way ANOVA followed by Tukey test. Bars represent the mean of three biological replicates, and the error bars represent the standard error of the mean.

We next investigated whether arabinofuranosidase (ABF) activity increased during wheat–*Z. tritici* interactions. With this aim, we extracted apoplastic fluid from leaves of four wheat cultivars (Setenil, Aubusson, Drifter, and Titlis) and quantified the total ABF activity, using the chromogenic substrate *p*-nitrophenyl-α-L-arabinofuranoside (O-PNPAF), in both non-infected (mock) and plants infected with *Z. tritici* (3D7-GFP strain) at 7 dpi (Fig. 1B). Our biochemical analysis showed a basal ABF activity in all non-infected cultivars, while *Z. tritici* presence led to a significant increase in ABF activity (Fig. 1B). We next determined if the enhanced ABF activity during wheat-*Z. tritici* interactions was due to endogenous wheat arabinofuranosidases or to *Zt*GH54. We first obtained a disruption mutant line of *ZtGH54* (Δ*Ztgh54*). Subsequently, we measured the apoplastic ABF activity in the cultivar Setenil infected with 3D7-GFP or Δ*Ztgh54*. Unlike the control strain 3D7-GFP, the Δ*Ztgh54* deletion mutant did not induce a significant increase in ABF activity, comparable to that of non-infected plants (Fig. 1C). Overall, the results indicate that the increased ABF activity detected during wheat infection is largely mediated by *Zt*GH54, suggesting this enzyme as the major source of pathogen-derived ABF activity within the apoplastic space.

### *Zt*GH54 is a *bona fide* α-L-arabinofuranosidase

To determine the biochemical properties and substrate specificity of *Zt*GH54, this protein was expressed in *Pichia pastoris*. In addition to the wild-type version of the enzyme (*Zt*GH54^Wt^), we obtained a mutant variant modifying the predicted catalytic residues, Glu218 and Asp293, by alanine (*Zt*GH54^E218A/D293A^). After two chromatographic steps, using cation-exchange and exclusion molar columns, the homogeneity of pure enzyme was determined by SDS-PAGE (Supp. Fig 1). In the case of the *Zt*GH54^E218A/D293A^ variant, a single protein band was also obtained with the same protocol, with an apparent molecular mass practically identical to that of the wild type enzyme (32 kDa). The identification of the *Zt*GH54 protein band, after SDS-PAGE, by nanosystem liquid chromatography-tandem mass spectrometry, confirmed that the protein is an arabinofuranosidase (table S1). The biochemical analysis of purified proteins, using O-PNPAF as substrate, demonstrated that *Zt*GH54^Wt^ exhibited ABF activity, whereas the mutant variant *Zt*GH54^E218A/D293A^, as expected, was inactive (Fig. 2A). The crude extract of the ascomycete *Talaromyves amestolkiae*, as a source of β-glucosidases and β-xylosidases (*42*, *43*) was used as a positive control, and 20 mM sodium acetate, pH 4.0 (mock) as negative control.

**FIGURE 2.**
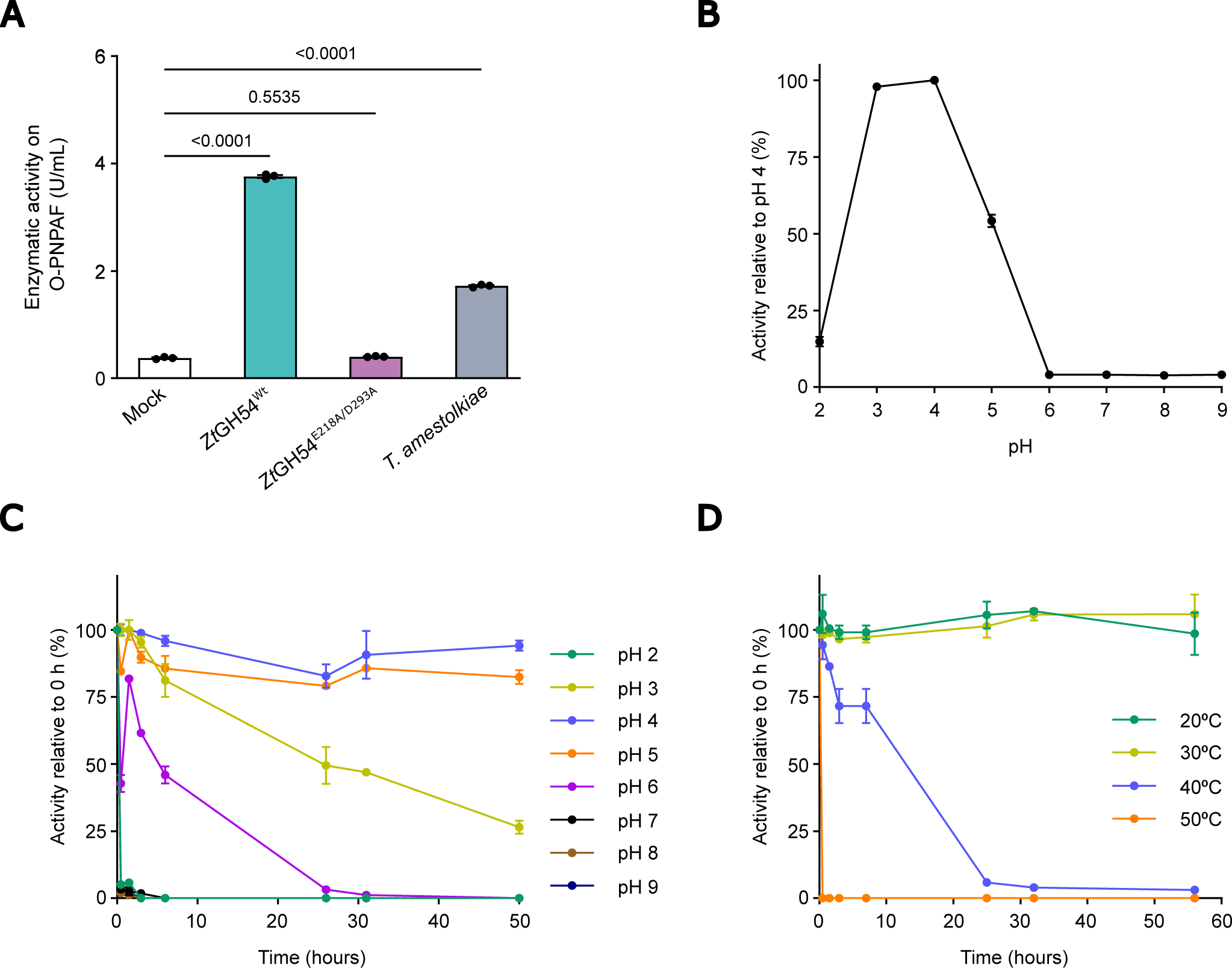
*Zt*GH54 is a stable *bona fide* α-L-arabinofuranosidase. **A.** Enzymatic activity of both purified recombinant *ZtGH54*^Wt^ and *Zt*GH54^E218A/D293A^ proteins on *p*-nitrophenyl-α-L-arabinofuranoside (O-PNPAF). The crude extract of *Talaromyces amestolkiae* and 20 mM sodium acetate at pH 4.0 (mock) were used as positive and negative controls, respectively. *P* values according to one-way ANOVA followed by Dunnett test are displayed in the plot. Bars represent the mean of three biological replicates, and the error bars represent the standard error of the mean. **B.** Optimum pH for enzymatic activity of *ZtGH54*^Wt^ on O-PNPAF in a pH range from 2 to 9 after 15 min incubation at 50 °C. 100% of enzymatic activity corresponded to the highest activity value observed (pH4). **C-D.** pH and thermal stability of *ZtGH54*^Wt^ on O-PNPAF in a pH range from 2 to 9 (**C**) or in a range of temperatures from 20 to 50 °C (**D**) over time. 100% corresponded to the activity value observed at 0 h. In B to D, dots represent the mean of two biological replicates and the error bars represent the standard deviation.

β-galactosidase (EC 3.2.1.23) and β-1,4-xylosidase (EC 3.2.1.37) activities of both *Zt*GH54 variants were also determined using *o*-nitrophenyl-β-D-galactopyranoside and *p*-nitrophenyl-β-D-xylopyranoside, respectively. Although these activities have been predicted in some GH54 family members (CAZy database), no detectable activity against these substrates was detected (Supp. Fig. 2A-B), suggesting that *Zt*GH54^Wt^ functions as a *bona fide* ABF. To complete the characterization of *Zt*GH54, its pH profile and long-term stability at different pH values and temperatures were determined, using O-PNPAF as substrate (Fig. 2B-D). The purified enzyme showed a narrow acidic activity profile with a maximum activity at pH 3.0 and 4.0. Stability assays across a pH range of 2.0 to 9.0 showed that *Zt*GH54 is highly stable under acidic conditions, retaining activity at pH 4.0 and 5.0 over 50 hours, whereas neutral and alkaline conditions triggered a progressive decline of catalytic capacity (Fig. 2B-C). Thermal stability assays performed at pH 4.0 showed that *Zt*GH54 remains highly active for up to 56 hours at 20 and 30 °C, while incubation at 50 °C caused a rapid and complete inactivation of ABF activity after 30 min (Fig. 2D). These results indicate that *Zt*GH54 is highly stable and active under acidic conditions of the host apoplastic space.

To resolve the structural basis underlying the substrate preference of *Zt*GH54 at the atomic level, the AlphaFold2-predicted structure of the enzyme (Supp. Fig. 3A) was subjected to molecular docking with arabinan using DiffDock, followed by 100-ns all-atom molecular dynamics (MD) simulations of complexes with arabinan, arabinoxylan, and glucoxylan (Supp. Fig. 3). MD simulations showed that arabinan and arabinoxylan remained stably bound to *Zt*GH54. In both complexes, a single α-1,3-linked arabinofuranosyl residue was consistently positioned within the catalytic pocket and remained stably engaged throughout the simulations. In contrast, glucuronoxylan rapidly dissociated from the active site (Supp. Fig. 3C). In the final MD conformations, the arabinofuranosyl residue of both arabinan and arabinoxylan was positioned within the catalytic pocket, establishing interactions with D216, a conserved substrate-binding residue in GH54 α-L-arabinofuranosidases (*44*), and with E218 and D293, two catalytic residues (Fig. 3A and B). Comparison of *Zt*GH54 with the crystal structure of *Aspergillus kawachii* GH54 bound to arabinose (PDB 1WD4) showed that the arabinofuranosyl moieties of both substrates adopt a highly similar orientation within the catalytic pocket (Supp. Fig. 3D; Table S2), supporting the model obtained during the MD simulations.

**FIGURE 3.**
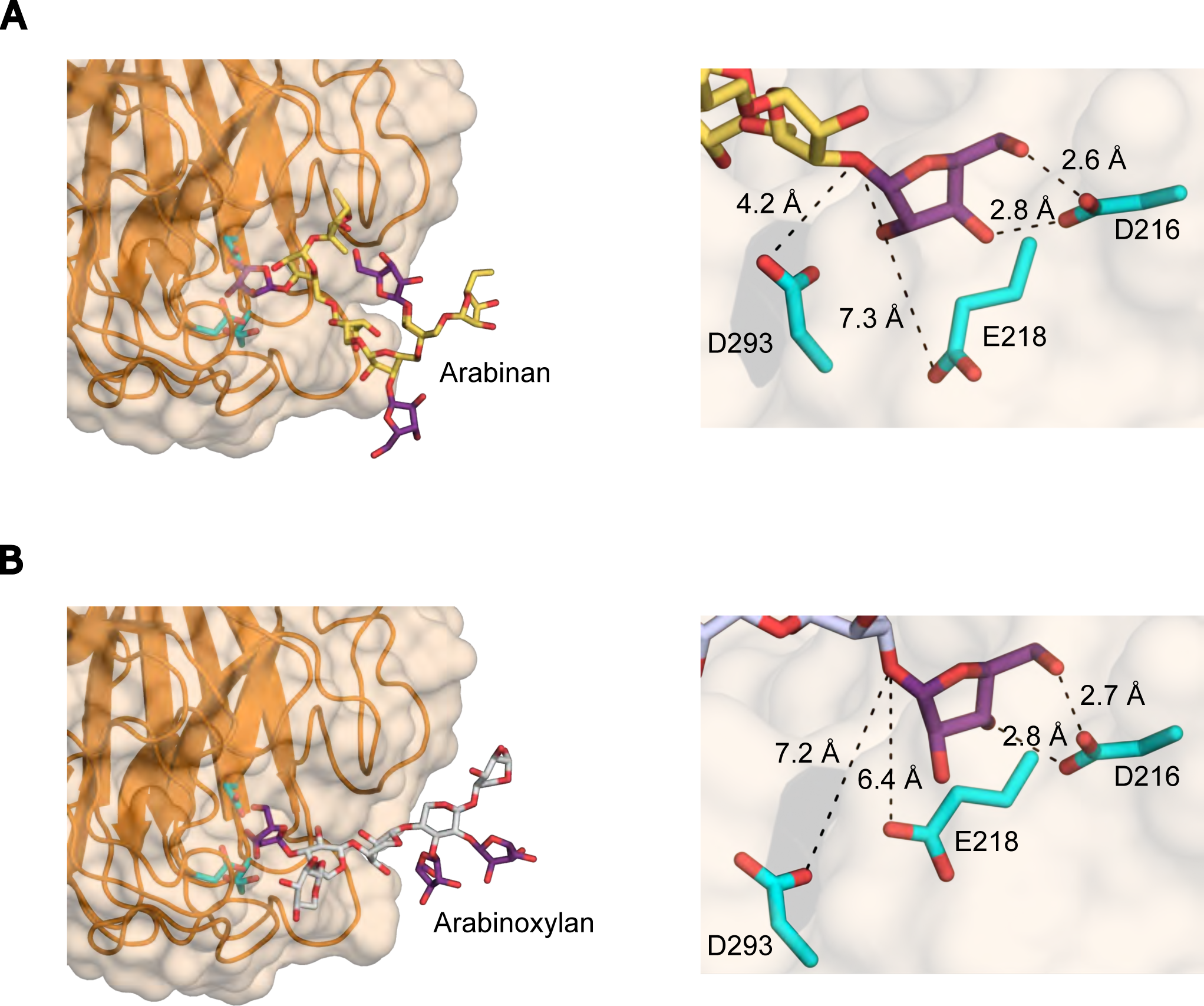
*Zt*GH54 binds arabinan and arabinoxylan. **A-B.** Final conformations obtained after 100-ns all-atom molecular dynamics simulations of *Zt*GH54 in complex with arabinan (**A**) and arabinoxylan (**B**). The main chains of arabinan and arabinoxylan are shown in yellow and grey, respectively, whereas the α-L-arabinofuranosyl side chains are shown in purple. The arabinan (**A**, right panel) and arabinoxylan (**B**, right panel) remained stably bound within the catalytic pocket throughout the simulations. The distances between the catalytic (E218 and D293) and the substrate-binding (D216) residues and the ligand are indicated.

### *Zt*GH54 supports *Z. tritici* growth

We hypothesized that the α-L-arabinofuranosidase activity of *Zt*GH54 could potentially contribute to nutrient acquisition from plant cell walls. We quantified the growth in the presence of arabinan or arabinoxylan of the control 3D7-GFP strain and the knockout line (Δ*Ztgh54*), along with a complementation line carrying the wild-type version of *Zt*GH54 (*ΔZtgh54/ZtGH54^Wt^*) and a complementation line expressing the catalytically inactive variant of *Zt*GH54 (*ΔZtgh54/ZtGH54^E218A/D293A^*). The strains cultured in the absence of the polysaccharides did not show growth differences (Supp. Fig. 4A). In contrast, the Δ*Ztgh54* mutant exhibited a significant reduction in biomass accumulation, compared to the control, when the medium was amended with arabinan or arabinoxylan (Fig. 4A-C). Genetic complementation of the mutant strain with *ZtGH54^Wt^*gene fully rescued this growth defect. Conversely, complementation with the catalytically inactive *ZtGH54^E218A/D293A^* gene did not restore the growth phenotype of the knockout line (Fig. 4A-C). These results indicate that arabinose, the *Zt*GH54 product, could be used as a nutrient by *Z. tritici*. To confirm this hypothesis, we grew the 3D7-GFP strain in Vogel’s minimal medium supplemented with arabinose (Fig. 4D). As expected, 3D7-GFP growth was minimal in the absence of any carbon source, but grew in the presence of arabinose, corroborating that *Z. tritici* possesses the necessary metabolic machinery to assimilate this monosaccharide. Finally, to rule out the possibility that any nutritional phenotypes resulted from impaired growth or altered stress tolerance, we evaluated the in vitro growth of the *Z. tritici* strains in yeast malt sucrose (YMS) medium in the presence of NaCl, H_2_O_2_, or 24 °C. The results indicate that the growth and development were similar among 3D7-GFP, Δ*Ztgh54*, and the complementation lines (Supp. Fig. 4B). Overall, these results indicated that *Zt*GH54 plays an important role in nutrient acquisition from plant cell walls, potentially during host colonization.

**FIGURE 4.**
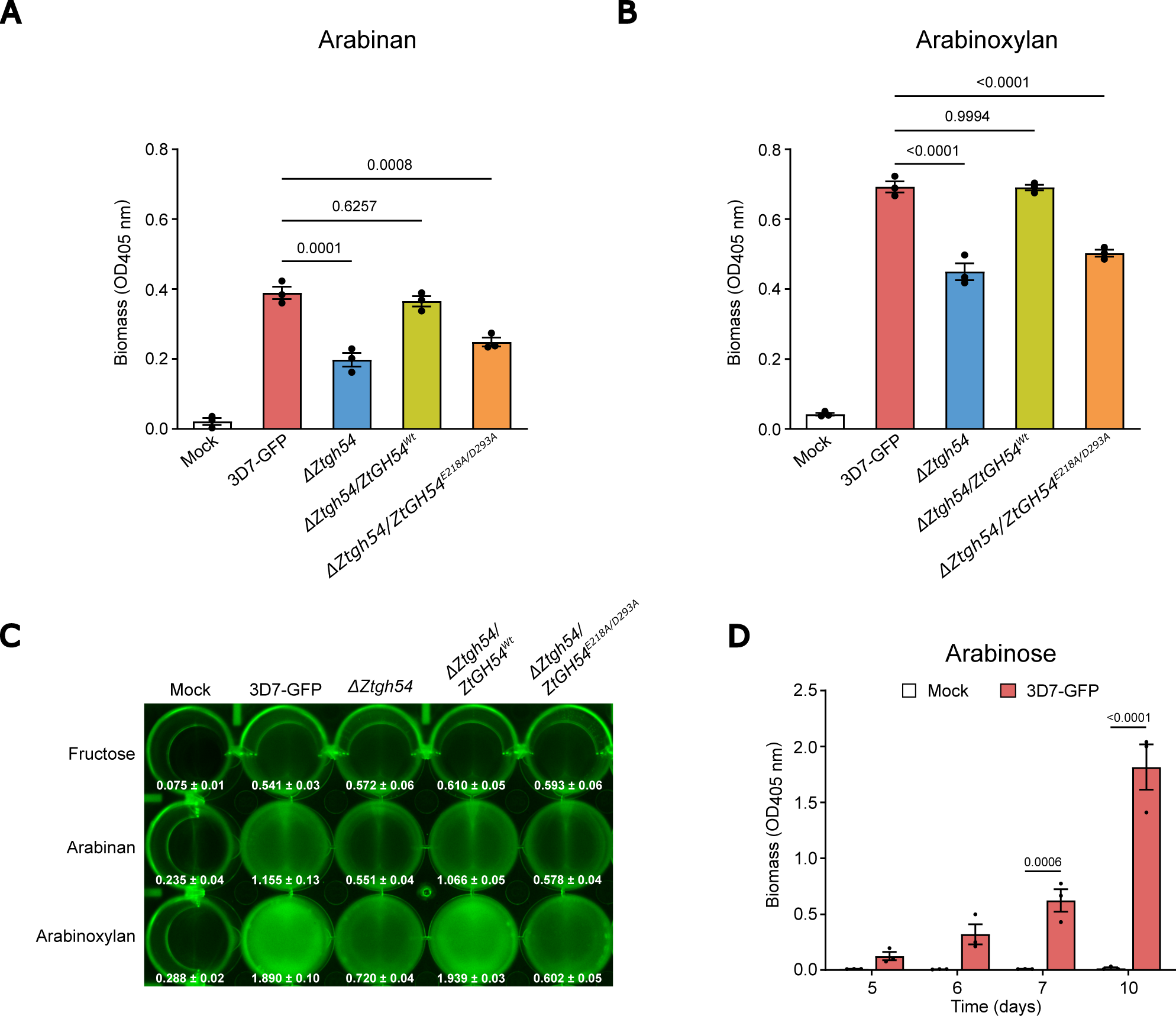
*Zt*GH54 supports *Z. tritici* growth in arabinan and arabinoxylan. **A-B.** OD_405nm_ measure as a proxy of biomass of 3D7-GFP, *ZtGH54* deletion mutant (*ΔZtgh54*), and complementation lines with *ZtGH54* wild-type version (*ΔZtgh54/ZtGH54^Wt^*) or *ZtGH54* with the mutated catalytic site (*ΔZtgh54/ZtGH54^E218A/D293A^*) grown at 18 °C for 5 days on liquid Vogel’s minimal media supplemented with either 0.5% w/v arabinan (**A**) or 0.5% w/v arabinoxylan (**B**). *P* values according to one-way ANOVA followed by Dunnett test are displayed in the plots. **C.** GFP fluorescence of 3D7-GFP, *ΔZtgh54*, *ΔZtgh54/ZtGH54^Wt^*, and *ΔZtgh54/ZtGH54^E218A/D293A^* strains grown at 18 °C for 5 days on liquid Vogel’s minimal media supplemented with either 0.5% w/v fructose (upper line), 0.5% w/v arabinan (middle line) or 0.5% w/v arabinoxylan (bottom line). The values of each well represent the mean of three biological replicates and the standard error of the mean. **D.** OD_405nm_ measure of non-inoculated (mock, white bar) and inoculated with 3D7-GFP (red bar) grown at 18 °C for 5 to 10 days on liquid Vogel’s minimal media supplemented with 0.5% w/v arabinose. *P* values according to two-way ANOVA followed by Šídák test between 3D7-GFP and mock treatment are displayed in the plot. Bars represent the mean of three biological replicates, and the error bars represent the standard error of the mean.

### *Zt*GH54 hydrolyzes the immunogenic DAMP XA^3^XX to suppress wheat defense

We next explored whether *Zt*GH54 could also hinder the accumulation of immunogenic arabinoxylan-derived oligosaccharides acting as DAMPs. Among these oligosaccharides, the pentasaccharide XA^3^XX (3^3^-α-L-arabinofuranosyl-xylotetraose) triggers defense responses in dicotyledonous plants (*37*, *38*) and could be hydrolyzed by *Zt*GH54 to release the arabinose residue. To explore whether XA^3^XX is a substrate of *Zt*GH54, we incubated both purified recombinant *Zt*GH54^Wt^ and the mutant *Zt*GH54^E218A/D293A^ proteins with commercial XA^3^XX oligosaccharide and quantified the released products using high-performance anion-exchange chromatography with pulsed amperometric detection (HPAEC-PAD). Our quantitative analysis showed that after three hours of incubation at 20 °C, the reaction mixtures containing the *Zt*GH54^Wt^ enzyme produced high concentrations of free arabinose and linear xylotetraose (Xyl4), comparable to those generated by the commercial positive control α-L-arabinofuranosidase from *Aspergillus niger* (E-AFASE) (Fig. 5A-B). In contrast, the incubation with the inactive *Zt*GH54^E218A/D293A^ protein did not hydrolyze the substrate, resulting in only minimal levels of arabinose and no detectable accumulation of Xyl4 (Fig. 5A). To confirm whether this enzymatic activity operates during wheat infection, we quantified xylotetraose accumulation in cultivar Setenil under non-infected (mock) conditions or infected with either the control strain 3D7-GFP or the deletion mutant of *ZtGH54* (Δ*Ztgh54*), using hydrophilic chromatography combined with electrospray ionization mass spectrometry (HILIC/ESI-MS). Leaves inoculated with the Δ*Ztgh54* strain showed a significantly lower content of xylotetraose compared to the control 3D7-GFP strain (Fig. 5B). These results highlight that *Zt*GH54 removes the branched α-L-arabinofuranosyl residue from XA^3^XX and actively contributes to the processing of AX-derived elicitors within the wheat apoplast during infection.

**FIGURE 5.**
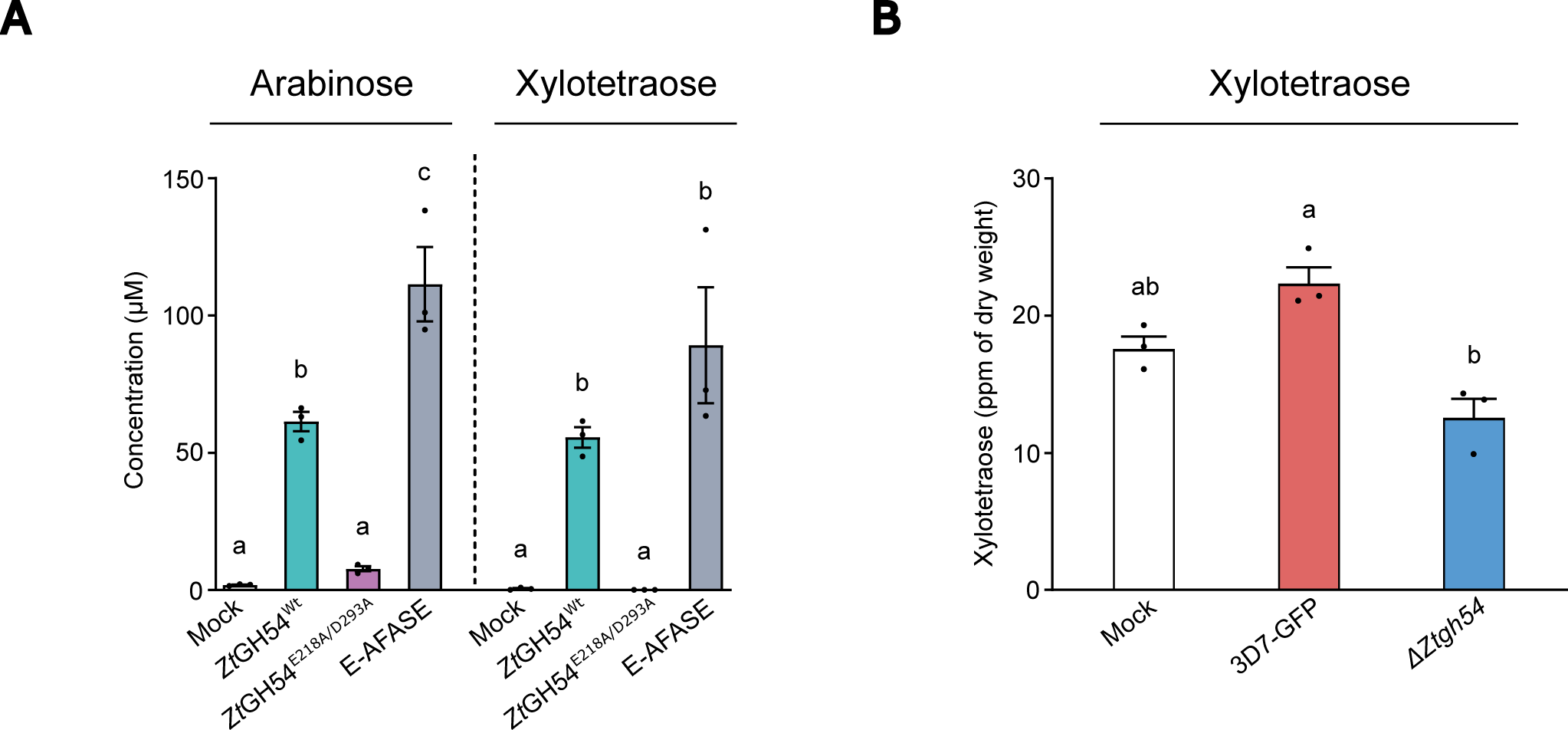
*Zt*GH54 hydrolyzes arabinoxylan-derived elicitors. **A.** Quantification by HPAEC-PAD of arabinose and xylotetraose upon incubation of both purified recombinant *Zt*GH54^Wt^ and *Zt*GH54^E218A/D293A^ proteins with XA^3^XX. Commercial α-L-arabinofuranosidase from *Aspergillus niger* (E-AFASE) and 20 mM sodium acetate (pH 4.0) buffer without enzymes (mock) were used as positive and negative controls, respectively. **B.** Quantification by HILIC/ESI-MS of xylotetraose released in cultivar Setenil non-infected (mock) and infected (7 days post infection) with 3D7-GFP or *ZtGH54* deletion mutant (*ΔZtgh54*). In A-B, different letters indicate significant differences (*P* value <0.05) according to one-way ANOVA followed by Tukey test. Bars represent the mean of three biological replicates, and the error bars represent the standard error of the mean.

Having established that *Zt*GH54 hydrolyzes XA^3^XX, we evaluated whether this enzymatic cleavage affects wheat immunity activation. Xylotetraose (Xyl4) triggers ROS production in wheat (*38*). However, the role of XA^3^XX as a DAMP in wheat has not been demonstrated. To determine whether this branched pentasaccharide activates host defense responses, we quantified the total ROS accumulation in leaves of Setenil, Aubusson, Drifter, and Titlis cultivars upon treatment with XA^3^XX. We treated leaf discs with either XA^3^XX or Xyl4 oligosaccharides, while water (mock) and chitohexaose (CHI6) were used as negative and positive controls, respectively. We observed that wheat leaves underwent a broad-spectrum oxidative burst upon XA^3^XX treatment, leading to significant ROS accumulation across all tested wheat cultivars compared to the mock control (Fig. 6A-B). In contrast, the response to the linearized Xyl4 product from *Zt*GH54 was cultivar-specific. Treatment with Xyl4 caused a significant increase in ROS accumulation only in Drifter and Titlis leaf tissues, while Setenil and Aubusson showed similar ROS levels to the mock treatment (Fig. 6A-B). These results indicate that the single α-L-arabinofuranosyl branch of XA^3^XX is essential for its recognition as a broad-spectrum DAMP in wheat.

**FIGURE 6.**
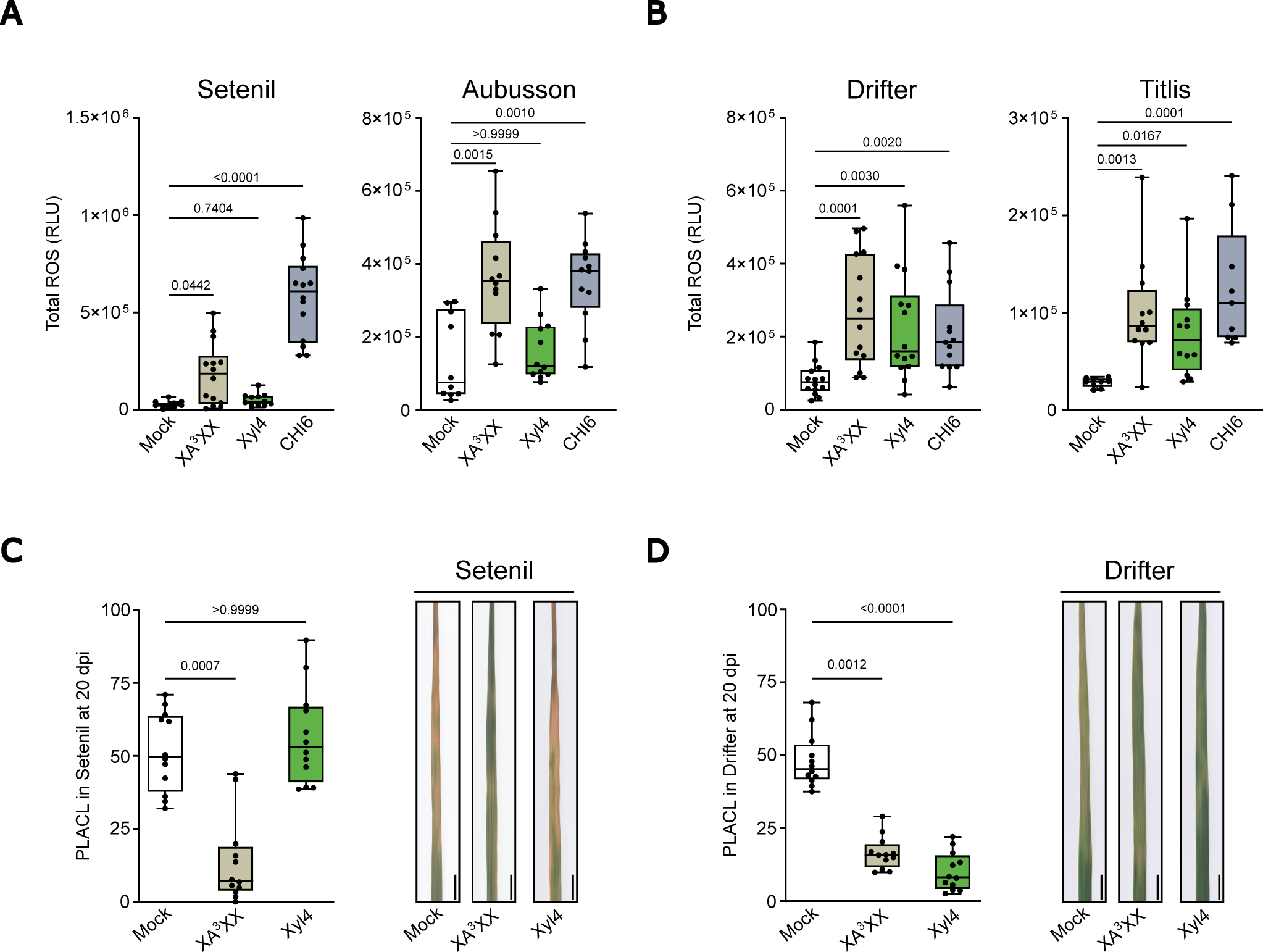
XA^3^XX triggers broad-spectrum ROS accumulation and resistance to *Z. tritici*, whereas xylotetraose responses are cultivar-specific. **A-B.** Total reactive oxygen species (ROS) production, estimated as cumulative relative luminescence units (RLUs) over 60 min, in response to 250 µM 3^3^-α-L-Arabinofuranosyl-xylotetraose (XA^3^XX) or 250 µM xylotetraose (Xyl4) in cultivar Setenil and Aubusson (**A**) or Drifter and Titlis (**B**). 100 μM hexaacetyl-chitohexaose (CHI6) and Milli-Q water (mock) were used as positive and negative controls, respectively. **C-D**. Virulence of *Z. tritici* (3D7-GFP strain), measured as percentage of leaf area covered by lesions (PLACL) at 20 days post infection (dpi), in cultivars Setenil (**C**) or Drifter (**D**) pretreated for 24 h with 0.1% v/v UEP-100 and 0.01% v/v Tween 20 (mock), 0.5 mM XA^3^XX or 0.5 mM Xyl4. Representative images of infected wheat leaves are displayed to the right of panels C and D. Scale bar, 1 cm. In A to D, *P* values according to Kruskal-Wallis test followed by Dunn test are displayed. In the boxplots, the middle line and the box represent the median and the interquartile range, respectively. Whiskers extend to the minimum and maximum values, and each black dot represents an individual datapoint.

To confirm whether Xyl4 and XA^3^XX protect wheat against infection, we performed *in planta* protection assays using one of the cultivars that responds to Xyl4, Drifter, and an additional one that did not respond to Xyl4, Setenil. We pretreated wheat leaves with XA^3^XX or Xyl4 24 hours before inoculating them with *Z. tritici*. XA^3^XX application to Setenil and Drifter leaves reduced symptom development compared to mock-treated plants, as quantified 20 dpi (Fig. 6C-D). In contrast, pretreatment with Xyl4 failed to protect Setenil plants (Fig. 6C), while Z. tritici lesions were reduced in Drifter (Fig. 6D). These results indicate that the branched α-L-arabinofuranosyl residue in XA^3^XX is crucial for protecting Setenil against *Z. tritici* infection. Overall, these findings reveal a mechanism of immune evasion at the apoplast where *Z. tritici* secretes *Zt*GH54 to hydrolyze the DAMP XA^3^XX. By converting this branched structure into a linear Xyl4 oligosaccharide, the pathogen deconstructs a broad-spectrum elicitor into a molecule whose immunogenic potential is restricted to only certain wheat cultivars.

### *Zt*GH54 contribution to *Z. tritici* virulence is wheat cultivar-dependent

To determine the contribution of *Zt*GH54 to *Z. tritici* virulence, infection assays were performed on the four previously tested wheat cultivars. In Setenil and Aubusson cultivars, the deletion of *ZtGH54* (Δ*Ztgh54*) resulted in a reduction in virulence, as demonstrated by significantly lower percentage of leaf area covered by lesions (PLACL) values, compared to the control 3D7-GFP strain (Fig. 7A). Ectopic complementation with the wild-type *ZtGH54* allele (*ΔZtgh54/ZtGH54^Wt^*) restored virulence, whereas expression of a catalytically inactive variant (*ΔZtgh54/ZtGH54^E218A/D293A^*) completely failed to rescue the knockout phenotype (Fig. 7A). These results suggest that the contribution of *Zt*GH54 to virulence in cultivars Setenil and Aubusson requires the catalytic activity. An opposite infection pattern was observed when evaluating Drifter and Titlis cultivars. In these cultivars, infection with Δ*Ztgh54* deletion mutant resulted in increased PLACL compared to 3D7-GFP. Complementation of Δ*Ztgh54* with *ZtGH54^Wt^* restored symptom levels to those observed in the control strain. The strain complemented with the catalytically inactive version also showed a reduction of disease symptoms compared to the knockout control (Fig. 7B), suggesting that the protein *Zt*GH54 might be recognized in these two cultivars. Taken together, these results indicate that *Zt*GH54 contributes to *Z. tritici* virulence in a host genotype-dependent manner.

**FIGURE 7.**
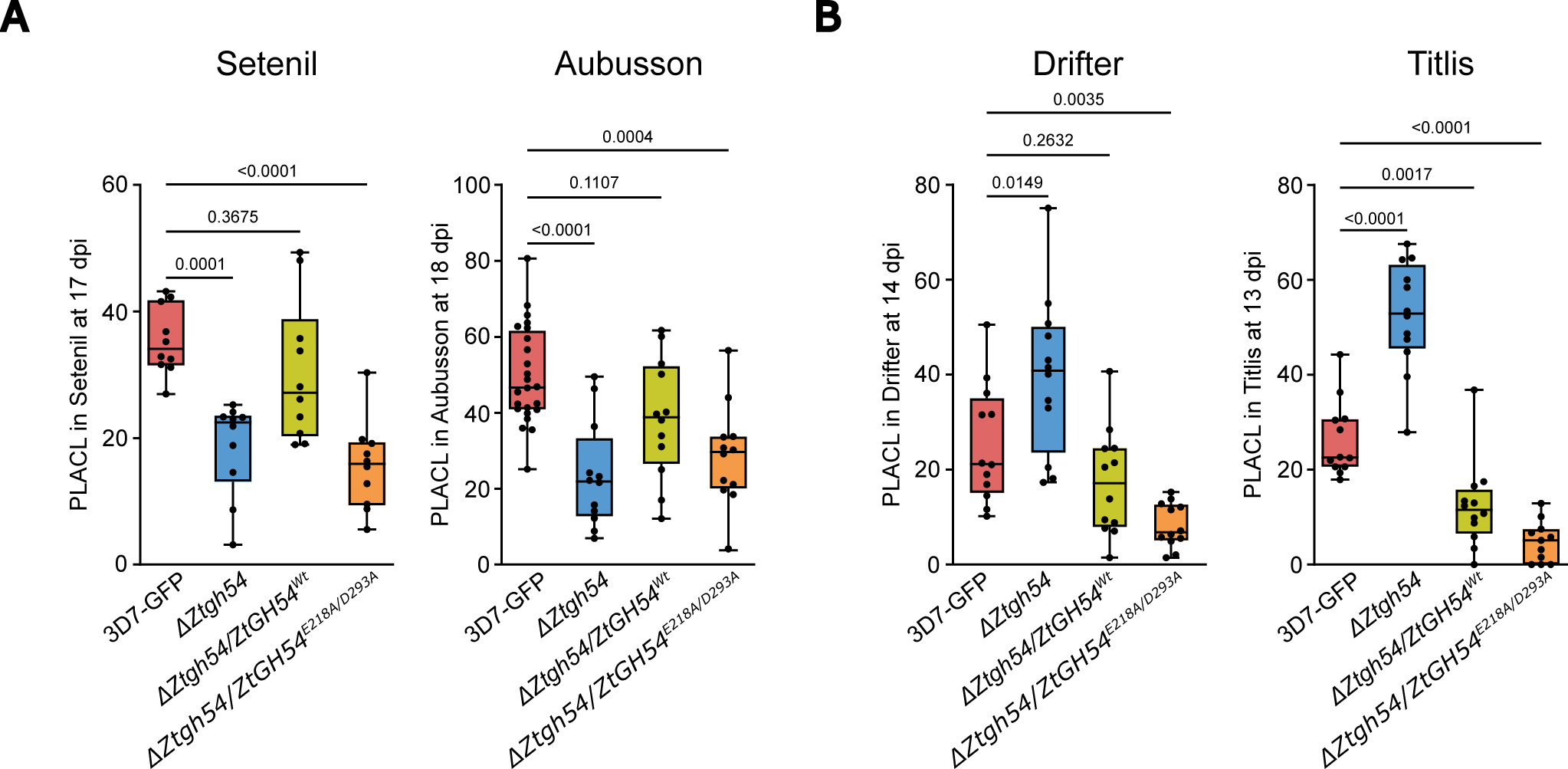
*Zt*GH54 contributes to *Z. tritici* virulence in a wheat cultivar-dependent manner. **A-B.** Percentage of leaf area covered by lesions (PLACL) produced by 3D7-GFP, *ZtGH54* deletion mutant (*ΔZtgh54*), and complementation lines with *ZtGH54* wild-type version (*ΔZtgh54/ZtGH54^Wt^*) or *ZtGH54* with the catalytic site mutated (*ΔZtgh54/ZtGH54^E218A/D293A^*) in cultivars Setenil and Aubusson (**A**) or Drifter and Titlis (**B**). *P* values according to one-way ANOVA followed by Dunnett test are displayed. In the boxplots, the middle line and the box represent the median and the interquartile range, respectively. Whiskers extend to the minimum and maximum values, and each black dot represents an individual data point.

## DISCUSSION

The plant-pathogen coevolutionary model has been largely described for the interaction between host resistance proteins and pathogen-secreted effectors (*45*, *46*). In this classical model, many effectors lack recognizable domains. However, other important effectors with hydrolytic domains, such as those involved in the modification or degradation of plant cell walls, also play an important role in determining the outcome of plant-pathogen interactions. These effectors, known as CWMEs (*47*), are secreted by pathogens during plant infection to acquire nutrients and facilitate host colonization (*8*, *10*). However, the enzymatic activity of CWMEs may come at a cost for pathogens, as cell wall degradation can release DAMPs that are perceived by the host and trigger immune responses. Therefore, pathogens need to balance plant cell wall degradation with the need to minimize host detection (*2*). A strategy used by the fungal pathogen *Z. tritici* to prevent early recognition by the wheat immune system is the delayed expression of the endoglucanase *ZtGH45*, thereby preventing the early release of immunogenic MLGs (*19*). Here, we describe a complementary mechanism, in which the α-L-arabinofuranosidase *Zt*GH54 hydrolyzes cell wall-derived elicitors.

Biochemical analysis demonstrated that *Zt*GH54 functions as an α-L-arabinofuranosidase, lacking β-galactosidase and β-1,4-xylosidase activity, and exhibits optimal activity under conditions encountered in the leaf apoplast (*48*). Molecular docking simulations analysis indicated that the catalytic residues E218 and D293, together with the substrate-binding residue D216, form a well-defined active-site pocket that accommodates α-L-arabinofuranosyl residues from arabinoxylan and arabinan. The activity of *Zt*GH54 releases L-arabinose, which can be utilized by *Z. tritici* as a carbon source, thereby linking *Zt*GH54 enzymatic activity to nutrient acquisition from plant cell wall components.

One of the substrates of *Zt*GH54 is arabinoxylan, a highly abundant wheat cell wall component. Specifically, it can use arabinoxylan-derived oligosaccharides as a substrate, such as 3^3^-α-L-arabinofuranosyl-xylotetraose (XA^3^XX), cleaving its glycosidic bond and releasing arabinose and xylotetraose (Xyl4). This enzymatic activity impacts apoplastic immune signaling in wheat. XA^3^XX is a DAMP capable of eliciting strong immune responses across diverse plant species, including Arabidopsis, tomato, and pepper, and conferring resistance against bacterial and fungal pathogens such as *P. syringae* and *S. sclerotiorum* (*37*, *38*). Consistently, XA^3^XX functions as a broad-spectrum elicitor in wheat, triggering immune responses across all tested cultivars. By removing the arabinose residue from XA^3^XX, *Zt*GH54 prevents the perception of the universal DAMP, producing linear Xyl4, which is recognized in other plant species (*38*, *39*), but in wheat its recognition is cultivar-specific (Fig. 8). Thus, by hydrolyzing a broadly recognized DAMP, *Zt*GH54 promotes evasion of host immunity.

**FIGURE 8.**
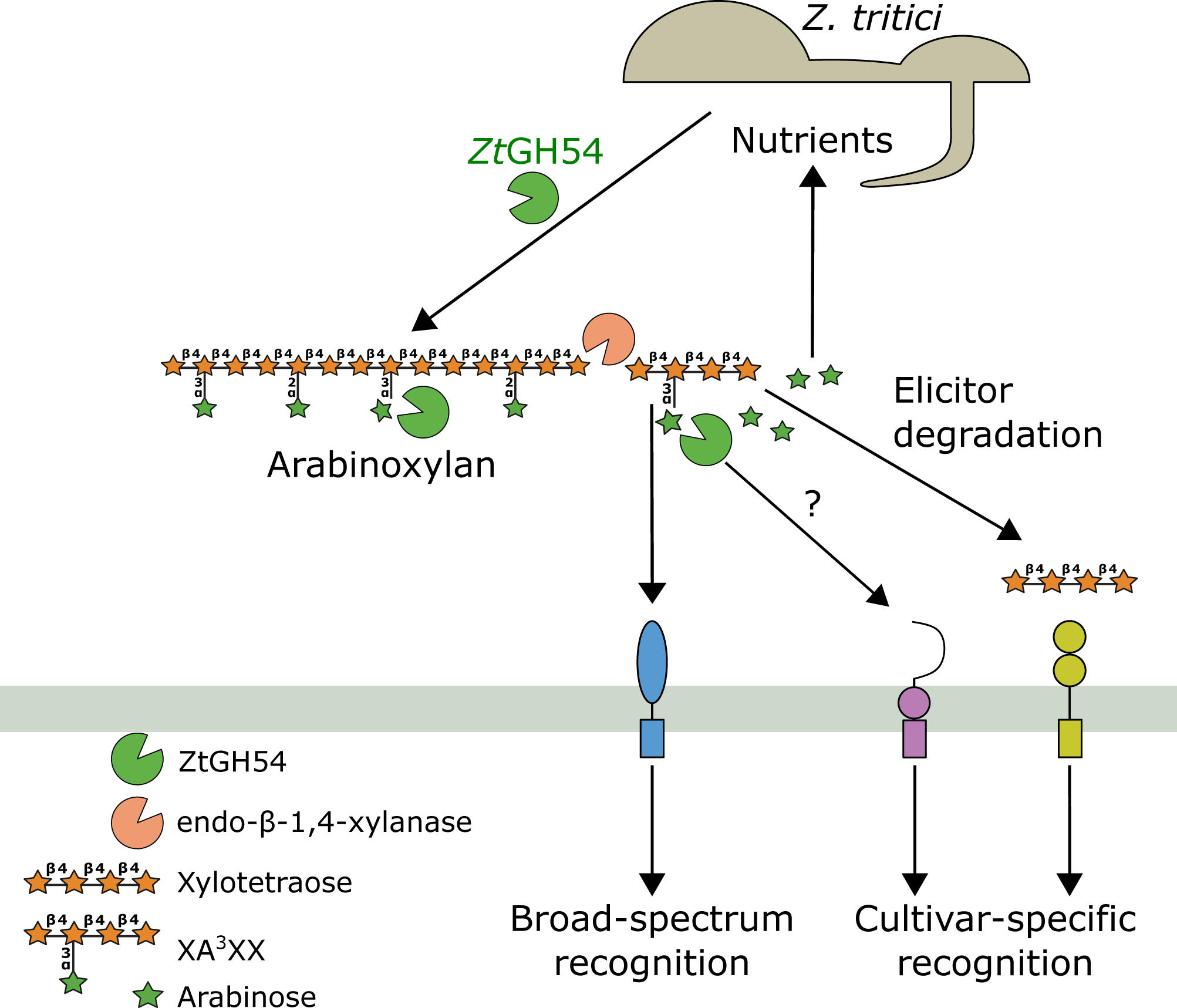
Proposed working model for the dual function of *Zt*GH54 during *Z. tritici* infection of wheat plants. At the early stages of infection, *Z. tritici* releases the α-L-arabinofuranosidase *Zt*GH54 into the apoplast of wheat, where it simultaneously plays a dual role in acquisition of nutrients and suppression of immune responses. *Zt*GH54 manipulates the apoplastic immune response by hydrolyzing the broad-spectrum elicitor XA^3^XX into xylotetraose, thus turning a general defense response into a cultivar-specific one.Furthermore, the *Zt*GH54 protein itself is recognized by specific wheat genotypes, triggering immune responses in a genotype-dependent manner.

According to the enzymatic activity of *Zt*GH54 and the cultivar-specific recognition of Xyl4, *Zt*GH54 functions as a key virulence factor in specific wheat cultivars such as Setenil and Aubusson. During infection with the control strain, ABF activity increases in the apoplast, accompanied by the accumulation of Xyl4 levels. In contrast, infection with the Δ*Ztgh54* disruption mutant did not result in an increase in ABF activity or Xyl4 accumulation. As a consequence, deletion of *ZtGH54* significantly reduces disease development in cultivars that recognize XA^3^XX. In contrast, in the cultivars Drifter and Titlis, the loss of function of *ZtGH54* does not impair virulence. This is consistent with the fact that these two cultivars can recognize both the product (Xyl4) and the substrate (XA^3^XX) of *Zt*GH54. Accordingly, in a recently published population genomics analysis, *Zt*GH54 was identified as a cultivar-specific virulence factor, which contributed to virulence in the wheat cultivar Aubusson (*36*). Remarkably, we observed that the knockout mutant in the cultivars was more virulent, and that this phenotype was complemented by expressing the wild-type and the catalytically inactive version of *Zt*GH54, indicating that these cultivars could additionally recognize the protein itself. The cultivar-specific contribution of *Zt*GH54 to virulence highlights host immune adaptation in wheat, suggesting that some cultivars have evolved specific surveillance systems to perceive the enzymatic *Zt*GH54 product, Xyl4, and/or the *Zt*GH54 protein itself. Wheat receptors recognizing Xyl4 and XA^3^XX remain to be identified. In *Arabidopsis thaliana*, perception of Xyl4 and XA^3^XX is mediated by IPG receptors (*38*). Remarkably, IGP homologues in wheat were identified in a genome-wide association study as a key locus associated with enhanced resistance to *Fusarium* crown rot and leaf rust (*49*). Potentially, this locus also contributes to the distinct perception of Xyl4 in wheat.

In conclusion, our findings establish the plant cell wall as a dynamic interface for the molecular arms race between plants and pathogens. Pathogens use enzymes such as *Zt*GH54 to hydrolyze broad-spectrum DAMPs like XA^3^XX, whereas plants counter this strategy by evolving receptors that can recognize either the enzymatic products or the CWMEs themselves (Fig. 8). This mechanism of host evasion complements the strategy employed by *Z. tritici* with the enzyme *Zt*GH45 (*19*), where the expression of the enzyme is delayed to the necrotrophic phase to delay the release of mixed-linkage glucan elicitors and avoid the early recognition by wheat. Together, these findings suggest that pathogens have evolved complementary strategies to evade host immunity, combining temporal regulation of CWMEs with enzymatic modification of immunogenic cell wall-derived oligosaccharides.

## MATERIALS AND METHODS

### *Z. tritici*, yeast and bacterial strains

The *Z. tritici* strain ST99CH_3D7 constitutively expressing the enhanced green fluorescent protein gene (3D7-GFP) (*50*, *51*) was used as control. *Escherichia coli* Stellar™ Competent Cells (Takara Bio Group, Japan) and *Agrobacterium tumefaciens* strain AGL1 were used for molecular cloning, plasmid propagation and *Z. tritici* transformation. *Pichia pastoris* X33 was used for protein expression.

### *Z. tritici* and bacterial culture conditions

*Z. tritici* was cultured in 100-mL Erlenmeyer flasks containing 50 mL of yeast sucrose broth (YSB; 1% w/v yeast extract and 1% w/v sucrose) or in yeast peptone dextrose (YPD; 1% w/v yeast extract, 2% w/v peptone, and 2% w/v dextrose), both supplemented with 50 μg/mL kanamycin sulphate. Cultures were maintained in the dark at 18 °C and 120 rpm for 6 days to obtain blastospores for the infection and developmental assays. To harvest the cells, cultures were filtered through sterile cheesecloth and centrifuged at 3,273 g for 15 min at 4 °C before being resuspended in Milli-Q water. Blastospore concentrations were quantified using a Neubauer counting chamber. *E. coli* was grown at 37 °C in LB medium (1.6% w/v tryptone, 1% w/v yeast extract, 0.5% w/v NaCl) supplemented with 50 μg/mL kanamycin sulphate. *A. tumefaciens* was cultured at 28 °C in LB medium containing 50 μg/mL kanamycin sulphate, 100 μg/mL carbenicillin and 50 μg/mL rifampicin.

### *Z. tritici* transformation

Targeted deletion of *ZtGH54* via homologous recombination was performed following the strategy previously described, using the In-Fusion HD Cloning Kit (Takara Bio Group, Japan (*18*). Briefly, approximately 1,000 bp upstream and downstream flanking regions of *ZtGH54* were PCR-amplified from 3D7 genomic DNA and assembled with the hygromycin resistance gene into the *KpnI*-and *SbfI*-linearized pCGEN plasmid. Complementation vectors for the wild-type and catalytic mutant (E218A and D293A) versions of *ZtGH54* included the CDS with introns, 993 bp upstream of the start codon and 725 bp downstream of the stop codon. The wild-type fragment was amplified from 3D7 genomic DNA, whereas the mutant variant was constructed by PCR-driven mutagenesis as described (*52*). Each insert was assembled into the *BamHI*-and *HindIII*-linearized pCGEN plasmid. All PCR amplifications were performed using Supreme NZYProof 2X Green Master Mix (NZYTech, Lda., Portugal) and the primers listed in table S3. *A. tumefaciens*-mediated *Z. tritici* transformation was performed and the transformants were selected on solid yeast malt sucrose broth (YMS; 0.4% w/v yeast extract, 0.4% w/v malt extract, 0.4% w/v sucrose, and 1.2% w/v Bacto agar) plates supplemented with 200 µg/mL cefotaxime and 25 µg/mL hygromycin or 100 µg/mL geneticin as previously described (*18*, *53*, *54*). The copy number of the inserts was estimated by qPCR, and at least three independent single-copy lines were selected and maintained for subsequent functional characterization.

### *Z. tritici* developmental and stress tolerance assays

Developmental and stress tolerance of the generated *Z. tritici* lines were evaluated by spotting a 3-µL drop of blastospores suspension (10^4^ spores/mL) onto YMS plates supplemented with 0.5 M NaCl, 1 M sorbitol, 1 mM H_2_O_2_, 430 mM Congo Red, or 200 ng/µL Calcofluor white. Plates were incubated at 18 °C in the dark for 7 days, except for an additional YMS plate that was incubated at 24 °C.

### Wheat infection and protection assays

The wheat (*Triticum aestivum* L.) cultivars Setenil, Aubusson, Drifter, and Titlis were grown in pots containing organic substrate under a long-day photoperiod (16 h of light at 18 °C/8 h of dark at 15 °C) and 60% relative humidity for 14-18 days, as described (*55*). The protection assay was performed following the strategy previously described (*19*), in which leaves were sprayed (1 mL/plant) 24 h before infection with a solution of 0.5 mM 3^3^-α-L-arabinofuranosyl-xylotetraose (O-XA3XX; Megazyme) or 0.5 mM xylotetraose (O-XTE; Megazyme) with UEP-100 (0.1% v/v; Croda) and Tween 20 (0.01% v/v) as adjuvants. A solution containing only the adjuvants was used as the mock control. For plant infections, blastospore suspensions at a concentration of 5·10^6^ spores/mL in 0.1% v/v Tween 20 were spray-inoculated on leaves (1 mL/plant). To maintain high humidity after inoculation, the plants were placed inside sealed plastic bags for 72 h. Disease symptoms on the second leaves were estimated between 13 and 18 dpi, as described (*56*). Percentage of leaf area covered by lesions (PLACL) and pycnidia density (pycnidia/cm^2^ of leaf or pycnidia/cm^2^ of lesion) were calculated using ImageJ (*57*) and an automated image analysis method (*58*).

### Apoplastic fluid extraction and ABF activity measurement

Apoplastic fluid (AF) was extracted from second leaves 7 dpi. Intact non-inoculated and inoculated leaves were rinsed in ice-cold Milli-Q water for 60 s and dried with absorbent paper. The leaves were cut into 5-cm segments using a sterile scalpel, and 14 segments were vertically aligned within a 50 mL conical tube with 40 mL of ice-cold 100 mM sodium acetate, pH 4.0 and subjected to three consecutive vacuum cycles of 5 min each, with abrupt depressurization between cycles. The segments were recovered, dried with absorbent paper, and transferred vertically to the barrel of a 20 mL syringe (without plunger), placed inside a new 50 mL conical tube, and centrifuged at 1,000 g for 15 min at 4 °C. The eluted AF was collected. To determine ABF enzymatic activity, 50 µL of AF was mixed with 200 µL of 7.4 mM *p*-nitrophenyl-α-L-arabinofuranoside (O-PNPAF; Megazyme) prepared in 100 mM sodium acetate, pH4.0. Reaction mixtures were incubated at 20 °C for 45 min at 1000 rpm, and the reaction was stopped by adding 500 µL of 70 mM Na_2_CO_3_. The absorbance of *p*-nitrophenol was measured at 410 nm using a quartz cuvette (Hellma, 108-002-10-40).

### Recombinant protein production and purification

The recombinant proteins *Zt*GH54^Wt^ and *Zt*GH54^E218A/D293A^, tagged with both c-myc epitope and a polyhistidine (6xHis) tag at the C-terminal was obtained using the *P. pastoris* heterologous expression system (Invitrogen; V195-20). The sequence of *ZtGH54^WT^* without its signal peptide was PCR-amplified using cDNA as a template obtained from RNA extracted from 3D7-GFP-infected (7 dpi) wheat plants. The mutant variant *ZtGH54^E218A/D293A^*was obtained by PCR-driven mutagenesis as described (*52*). All PCR amplicons were obtained using Supreme NZYProof 2X Green Master Mix (NZYTech, Lda., Portugal) and the primers listed in Table S3, and assembled into the *PstI*-and *XbaI*-linearized pPICZα B expression vector (Invitrogen; V195-20) using the In-Fusion HD Cloning Kit (Takara Bio Group, Japan). Upon transformation of *P. pastoris* by electroporation, transformants were selected on YPD supplemented with 50 μg/mL zeocin. For large-scale protein production, a preculture grown in Buffered Glycerol-complex Medium (BMGY; 1% w/v yeast extract, 2% w/v peptone, 100 mM potassium phosphate buffer (pH 6.0), 1.34% w/v yeast nitrogen base with ammonium sulfate, 4×10⁻⁵% w/v biotin, and 1% v/v glycerol) supplemented with 50 µg/mL zeocin was collected by centrifugation at 3,000 g for 5 min at 4 °C and then resuspended to an OD_600nm_ of 0.1 in Buffered Methanol-complex Medium (BMMY; 1% w/v yeast extract, 2% w/v peptone, 100 mM potassium phosphate buffer (pH 6.0), 1.34% w/v yeast nitrogen base with ammonium sulfate, 4×10⁻⁵% w/v biotin, and 0.5% v/v methanol). Cultures were grown at 28 °C and 200 rpm. Absolute methanol was added every 24 hours during the 7-day induction period. Upon centrifugation of the cells, protein production was verified by Western Blot using mouse monoclonal anti-c-Myc Tag antibody (Merck, 05-724-25UG) and the horseradish peroxidase (HRP)-conjugated anti-mouse secondary antibody (Sigma-Aldrich, A9044-2ML). Chemiluminescent signals were detected using the Pierce ECL Western Blotting Substrate kit (Thermo Scientific, 32209) and acquired with the iBright FL1000 imaging system (Invitrogen). Supernatants were sequentially clarified and concentrated by ultrafiltration through 5 kDa molecular weight cut-off membranes and dialyzed against 20 mM sodium acetate, pH 5.0. Recombinant *Zt*GH54 variants were purified by Fast Protein Liquid Chromatography using an ÄKTA Purifier system (GE Healthcare) through a consecutive two-step chromatographic procedure. The samples were first loaded into a HiTrap QFF column (Cytivia) equilibrated with 20 mM sodium acetate, pH 5.0. The target protein variants were recovered directly in the non-binding flow-through, while the majority of host proteins were retained in the column. The flow-through fractions with enzyme activity were combined and subsequently applied to a TSKgel G3000SWXL size-exclusion column of 5 µm (Tosoh Biosciences) equilibrated and eluted with 20 mM sodium acetate supplemented with 150 mM NaCl, pH 5.0. The sample was dialyzed against 20 mM sodium acetate, pH 4.0. In the case of the cultures of the *Zt*GH54^E218A/D293A^ variant, the enzyme was purified following the same protocol.

### Identification of the purified *Zt*GH54 protein

The purified protein was separated by SDS-PAGE on 10% polyacrylamide gel and stained using the Colloidal blue staining kit (Invitrogen). The protein band was cut into small pieces and digested overnight at 37 °C with 12.5 ng/mL porcine trypsin in 50 mM ammonium bicarbonate. Peptides were extracted using 100% acetonitrile and 0.5% v/v trifluoroacetic acid, purified using an OMIX C18 100 µL column (Agilent Technologies), and dried as previously described (*59*). The reconstituted sample was analyzed by nanosystem liquid chromatography-tandem mass spectrometry (nLC–MS/MS). The peptide separations were performed on the Vanquish Neo nano system (Thermo Scientific), and the MS analysis was performed using an QExactive mass spectrometer (Thermo Scientific), selecting the 10 most intense precursor ions for MS/MS fragmentation. The nLC–MS/MS data were analyzed with Proteome Discoverer software (Thermo Scientific, version 3.0.0.757) using the SEQUEST HT search engine. The search was performed against a *P. pastoris* in-house database (5650 total sequences) supplemented with the target protein sequence. The identified peptides were filtered using the PSM Validator node.

### Characterization of *Zt*GH54 variants in *P. pastoris*

Substrate specificity of purified recombinant proteins, *Zt*GH54^WT^ and *Zt*GH54^E218A/D293A^ variants, was analyzed against three substrates, *p*-nitrophenyl-α-L-arabinofuranoside (O-PNPAF; Megazyme), *p*-nitrophenyl-β-D-xylopyranoside (O-PNPX; Megazyme), and *o*-nitrophenyl-β-D-galactopyranoside (ONPG; Thermo Scientific). The reaction mixture consisted of 7.4 mM of *p*-nitrophenyl substrates in 100 mM sodium acetate, pH 4.0, at 50 °C and 1000 rpm (15 min for O-PNPAF and 30 min for O-PNPX and ONPG), in a total volume of 250 µL with 200 µL of substrate and 50 µL of 2 µM purified protein. The reactions were stopped by adding 500 µL of 70 mM Na_2_CO_3,_ and the released nitrophenol was measured spectrophotometrically (ε_410_ = 15,200 M^-1^cm^-1^). One unit of activity was defined as the amount of enzyme that hydrolyzes 1 μmol of nitrophenyl-derived substrate per minute.

The pH stability profile of *Zt*GH54 activity was determined by mixing 5 µM of protein in 100 mM Britton and Robinson buffer, pH range of 2-9, at 25 °C, for 50 h and constant agitation at 1000 rpm. Thermal stability was assessed in the same buffer (pH 4.0), at 20, 30, 40 and 50 °C, for 56 h under agitation at 1000 rpm. Aliquots of each sample (50 µL) were taken at 0, 0.5, 1.5, 3, 7, 25, 32, and 56 h and transferred to fresh tubes. In all biochemical characterization assays, activity to O-PNPAF was measured as described above.

### Structural modeling of *Zt*GH54 and molecular docking

The structure of *Zt*GH54, without the 22-aminoacid signal peptide, was predicted using AlphaFold2 through ColabFold (*60*). The highest-ranked model was selected for subsequent analyses and exhibited pLDDT values above 90 across the entire sequence. Structural models of arabinan, arabinoxylan, and glucuronoxylan were generated using the Glycan Modeler tool available in CHARMM-GUI (*61*). Three independent molecular docking runs with arabinan were carried out using different random seeds, generating 20 binding poses per run under default inference settings using DiffDock v1.1.3 (*62*). Predicted poses were ranked according to the DiffDock confidence score, and the highest-confidence arabinan pose was selected for further analyses.

### All-atom molecular dynamic simulations

System parameterization was performed using CHARMM-GUI (*61*) with the CHARMM36 force field (*63*). Each complex was solvated in a rectangular TIP3P water box with a 14 Å padding distance. Sodium and chloride ions were added to neutralize the systems and achieve a final salt concentration of 0.15 M. Water molecules and ions were incorporated using VMD (*64*). Long-range electrostatic interactions were treated using the Particle-mesh Ewald (PME) method (*65*). Short-range nonbonded interactions were smoothly switched between 8 and 10 Å, with a 12 Å cutoff used for pair-list generation. Energy minimization was carried out for 5,000 conjugate-gradient steps, followed by a 100 ps equilibration phase at 298 K and 1 atm. Three independent 100-ns production simulations were performed for each complex in the NPT ensemble at 298 K and 1 atm using the SHAKE algorithm (*66*). Temperature and pressure were controlled using Langevin dynamics and the Langevin piston method, respectively. A time step of 2 fs was employed, and trajectory coordinates were saved every 50,000 steps. Simulations were performed on the CBGP computing cluster using NAMD 2.14 multicore-CUDA on Tesla V100 GPUs. Trajectories were visualized and analyzed using VMD (*64*).

### Nutrition assay

Vogel’s minimal medium (Vogel_MGB, 1956) supplemented with 0.5% (w/v) L-arabinose (Thermo Scientific) as the sole carbon source, or 0.5% (w/v) D-fructose (Thermo Scientific) amended with either 0.5% (w/v) arabinan (P-ARAB; Megazyme) or 0.5% (w/v) arabinoxylan (P-WAXYL; Megazyme) was inoculated with a spore suspension of each *Z. tritici* line at a final concentration of 4·10^5^ spores/mL. Cultures were incubated in the dark at 18 °C and 120 rpm for 5 days. *Z. tritici* biomass was assessed by measuring the optical density of each culture at 405 nm (OD_405nm_) using a quartz cuvette (Hellma, 108-002-10-40), as described (*67*, *68*), and by quantifying the GFP fluorescence using a Varioskan Lux plate reader (Thermo Scientific) with excitation and emission wavelengths set at 489 nm and 515 nm, respectively.

### Oligosaccharide hydrolysis assays

To identify and quantify the released oligosaccharides, we followed a similar protocol as the one described (*19*). Enzymatic hydrolysis assays were performed in a total volume of 200 µL containing 3 µM purified protein and 250 µM O-XA3XX (Megazyme) in 50 mM sodium acetate, pH 4.0. Incubations were performed in a thermoshaker at 20 °C for 3 h with constant agitation at 1000 rpm. The reaction mixtures were analysed by high-performance anion-exchange chromatography with pulsed amperometric detection (HPAEC-PAD) using a 930 Compact IC Flex chromatography system equipped with a FlexiPAD pulsed amperometric detector (Metrohm). Oligosaccharides were separated at 40 °C on a Metrosep Carb 2 250/4.0 analytical column and a Metrosep Carb 2 Guard/4.0 guard column (Metrohm). Isocratic elution was performed using 200 mM NaOH and 150 mM sodium acetate as a mobile phase with a flow rate gradient distributed as follows: 0-12 min at 0.7 mL/min, 12.1-22.0 min at 0.9 mL/min, and 22.1-27.0 min at 0.7 mL/min. Oligosaccharide quantification was determined utilizing standard calibration curves of XA^3^XX (O-XA3XX; Megazyme), xylotetraose (O-XTE; Megazyme), and L-arabinose (Thermo Scientific). The α-L-arabinofuranosidase (E-AFASE; Megazyme) was used as positive control.

### Quantification of xylotetraose *in planta*

For oligosaccharide extraction, 10-15 mg of second leaves from infected plants harvested at 7 dpi were homogenized in liquid nitrogen, resuspended in 0.5 mL of deionized water, ultrasonicated for 10 minutes, and heated at 95 °C for 10 minutes. After cooling, samples were centrifuged for 10 minutes at 17,000 g. Supernatants were filtered through an Ultrafree-MC centrifugal filter 0.22 µm. Pellets were washed with deionized water, centrifuged, and the supernatant filtered and pulled together. Oligosaccharide-containing filtered supernatants were freeze-dried and resuspended in 1 mM ammonium formate. For hydrophilic chromatography combined with electrospray ionization mass spectrometry (HILIC/ESI-MS), samples were injected onto a XBridge Amide column (3.5 µm, 2.1 x 150 mm; Waters, MA, USA) with a flow of 0.5 mL/min. The mobile phase consisted of 1 mM ammonium formate in water (Eluent A) and 1 mM ammonium formate in 90% acetonitrile (Eluent B). The eluent program was as follows: 100 % B at 0 min, 72.2 % B at 12 min, 61.1 % B at 13-14 min, 0 % B at 15 min. A standard calibration was performed using xylotetraose (O-XTE; Megazyme), and quantification was based on the (545.1609; 581.1404; 591.1703) ± 0.005 m/z ions ([M-H]^-^; [M+Cl]^-^; [M+CHOO]^-^). The chromatograms and spectra were processed using Compass DataAnalysis (version 5.3; Bruker Daltonics).

### Reactive oxygen species (ROS) quantification

ROS accumulation was monitored using 12.6 mm^2^ discs collected from the second leaves of 14-day-old wheat plants as previously described (*19*). Oxidative burst assays were triggered by adding 50 µL of either 750 µM O-XA3XX (Megazyme), 750 µM O-XTE (Megazyme), 100 µM hexaacetyl-chitohexaose (O-CHI6; Megazyme), or a Milli-Q water control to each disc. Each experimental run evaluated at least eight leaf discs per treatment, and the experiment was conducted three independent times.

### Statistical analysis

Data were processed, analysed and visualized using Prism 9 software (GraphPad Software, San Diego, CA, USA). Outliers were identified and removed using the ROUT method (Q=1%). Data distributions were evaluated for normality utilizing the Shapiro-Wilk and Kolmogorov-Smirnov tests. For comparisons between two independent groups, two-tailed unpaired Student’s *t*-tests were applied. Normally distributed multiple groups were subjected to parametric ordinary one-way or two-way analysis of variance (ANOVA), followed by Dunnett’s or Šídák’s post hoc multiple comparisons tests, respectively. In contrast, multi-group datasets displaying non-normal distribution were analyzed by the non-parametric Kruskal-Wallis test coupled with Dunn’s test. Specific statistical tests, replicate number, and significance thresholds are indicated in the figure legends.

## Acknowledgments

This work was supported by grant PID2023-150977NB-I00 funded by MICIU/AEI/10.13039/501100011033 and by FEDER, UE and RYC2018-025530-I grant from the Spanish Ministry of Science, Innovation, and Universities to ASV; by the Severo Ochoa Program for Centres of Excellence in R&D (grant CEX2020-000999-S (2022–2025)) funded by MCIU/AEI/10.13039/501100011033 to AM; and by PID2023-150378OB-I00 funded by MCIN/AEI/10.13039/501100011033 to HM. SLC was supported by fellowship PREP2023-002003, and GVP by fellowship PRE2021-100446, both funded by MICIU/AEI/10.13039/501100011033 and FSE+. We thank Gero Steinberg and Sreedhar Kilaru (Exeter University, UK) for providing us with the 3D7-GFP strain, Cécile Lorrain (ETH Zurich, Switzerland) for providing us biological material and very helpful discussion. The Proteomics and Genomics Facility at the Centro de Investigaciones Biológicas Margarita Salas performed the protein identification. We are also grateful to Ignacio Solís (Agrovegetal S.A.) and Colin Dervillez (Limagrain) for providing us with Setenil and Aubusson seeds, respectively.

## Author contributions

CCL and ASV designed the study. CCL, DR, FdS, SLC, IVL, and GVP performed the lab work. CCL, DR, FdS, SLC, GVP, MGA, MJM, HM, and ASV analyzed the data and conducted statistical analyses. DR and HM designed, performed, and analyzed HPAEC-PAD and HILIC/ESI-MS results. FdS and MJM designed, performed, and analyzed the protein purification and biochemical characterization results. GVP and MGA designed, performed, and analyzed molecular docking and molecular dynamics simulation results. CCL and ASV wrote the manuscript. MGA, MJM, HM, AM, and ASV secured funding for the project. All the authors critically reviewed and edited the manuscript.

## Data and materials availability

All data needed to evaluate the conclusions in the paper are present in the paper and/or the Supplementary Materials.

