## Supplementary figures and images for "A pathogen α-L-arabinofuranosidase degrades host-derived immunogenic oligosaccharides to suppress plant immunity"

### Supplementary figures

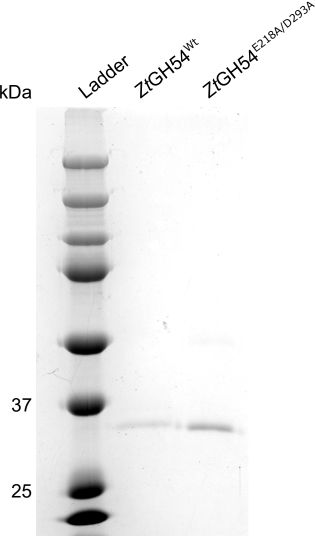

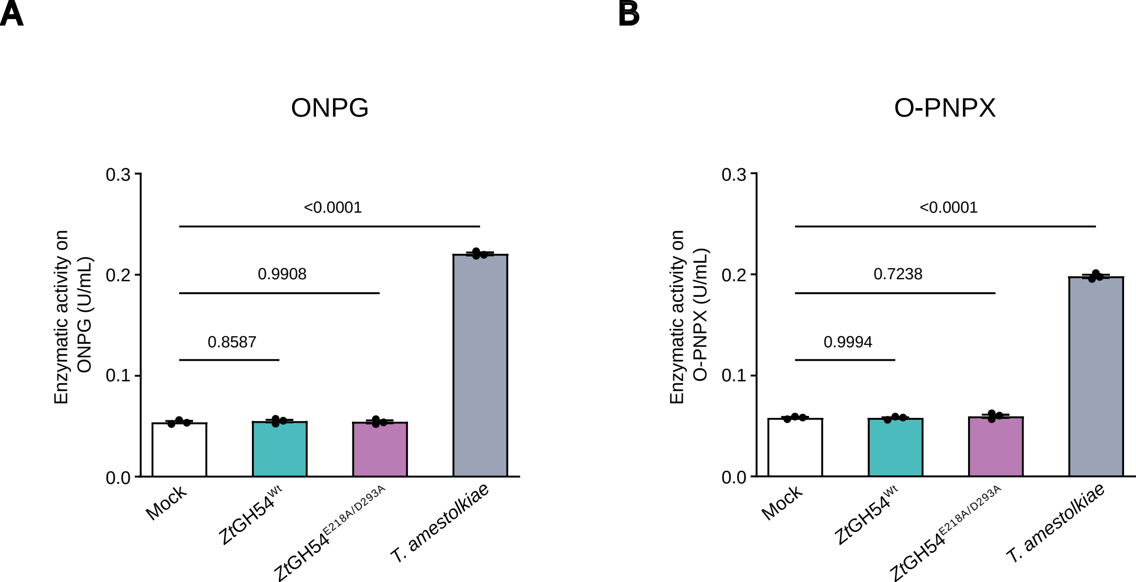

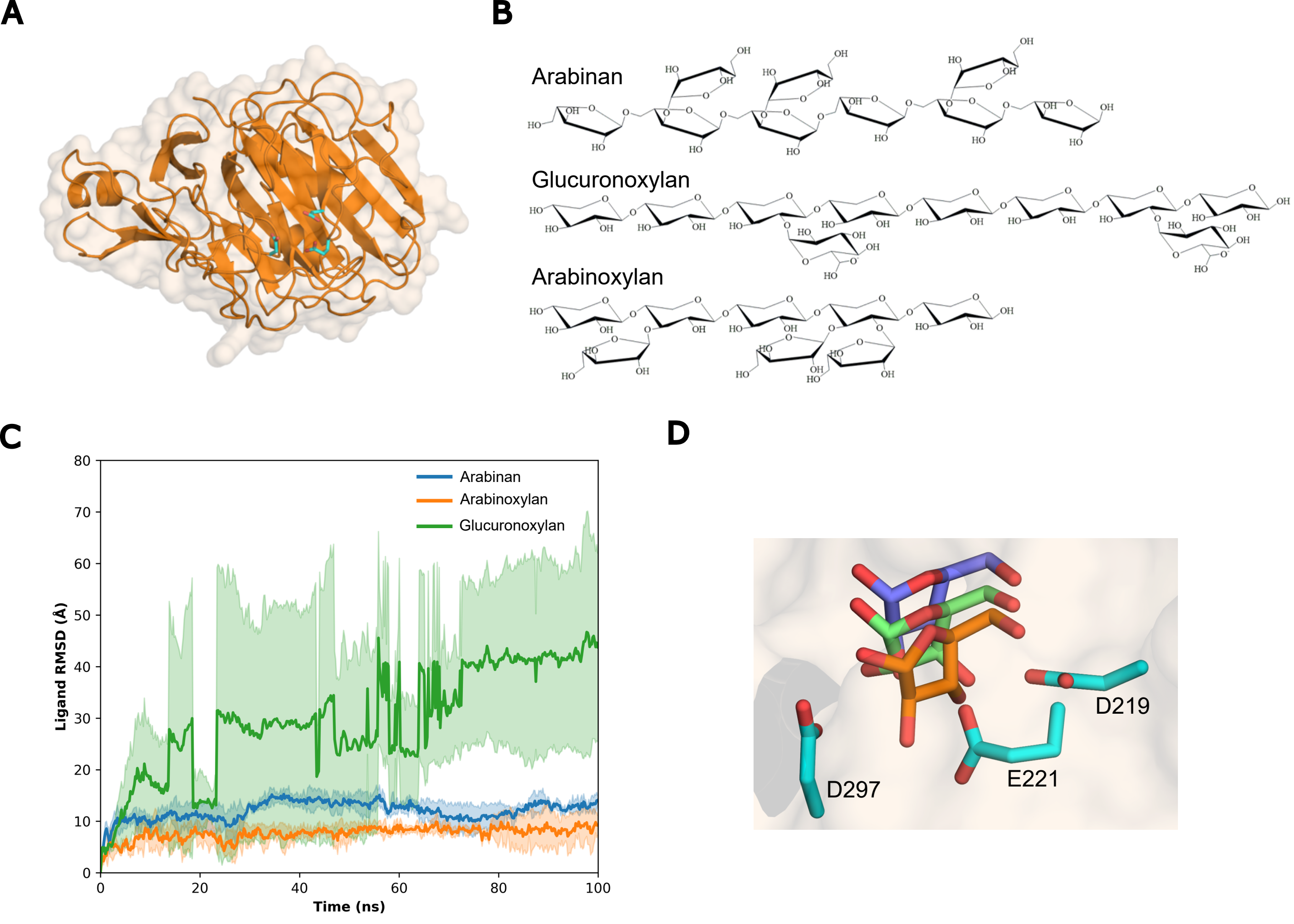

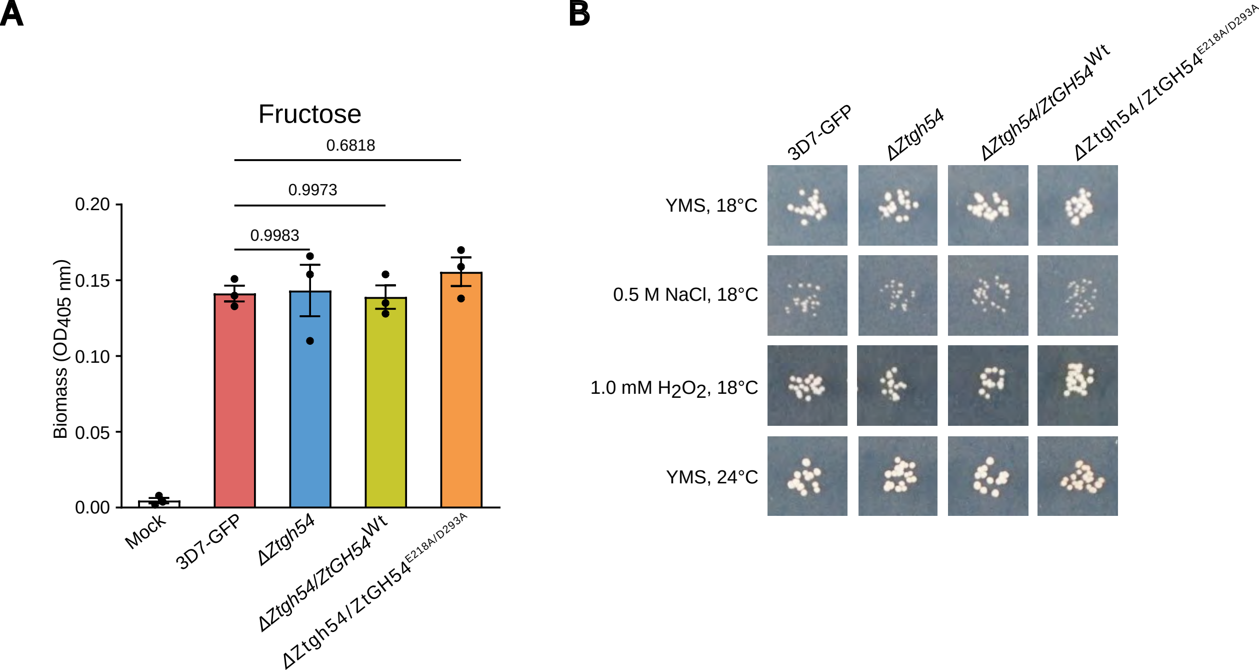
